# OBIA: An Open Biomedical Imaging Archive

**DOI:** 10.1101/2023.09.12.557281

**Authors:** Enhui Jin, Dongli Zhao, Gangao Wu, Junwei Zhu, Zhonghuang Wang, Zhiyao Wei, Sisi Zhang, Anke Wang, Bixia Tang, Xu Chen, Yanling Sun, Zhe Zhang, Wenming Zhao, Yuanguang Meng

## Abstract

With the development of artificial intelligence (AI) technologies, biomedical imaging data play an important role in scientific research and clinical application, but the available resources are limited. Here we present Open Biomedical Imaging Archive (OBIA), a repository for archiving biomedical imaging and related clinical data. OBIA adopts five data objects (Collection, Individual, Study, Series, and Image) for data organization, accepts the submission of biomedical images of multiple modalities, organs, and diseases. In order to protect personal privacy, OBIA has formulated a unified de-identification and quality control process. In addition, OBIA provides friendly and intuitive web interface for data submission, browsing and retrieval, as well as image retrieval. As of September 2023, OBIA has housed data for a total of 937 individuals, 4136 studies, 24,701 series, and 1,938,309 images covering 9 modalities and 30 anatomical sites. Collectively, OBIA provides a reliable platform for biomedical imaging data management and offers free open access to all publicly available data to support research activities throughout the world. OBIA can be accessed at https://ngdc.cncb.ac.cn/obia.

## Introduction

The introduction of advanced imaging technologies has greatly facilitated the development of non-invasive diagnoses. Currently, biomedical images can clearly depict the internal structure (anatomy), morphology, and physiological functions from the molecular scale to cellular, organ, tissue, lesion, and even the entire organism [1], providing crucial evidence for diagnosis and treatment response assessment [2,3]. Imaging data generated during patient visits has formed a huge accumulation. However, incomplete sharing systems make it challenging for researchers and clinicians to collaborate on utilizing these images to gain significant insights into health and disease [4]. Furthermore, the demand for rapid diagnosis promotes the application of artificial intelligence (AI) in biomedical imaging, and the development of reliable and robust AI algorithms requires sufficiently large and representative image datasets [5,6]. Thus, high-quality biomedical imaging data sharing plays an important role in promoting scientific discoveries and improving diagnostic accuracy.

The National Institutes of Health (NIH) in America sponsors several repositories. The Medical Imaging and Data Resource Center (MIDRC) [7] serves as an open-access platform for COVID-19-related medical images and associated data. Image and Data Archive (IDA) [8], NITRC Image Repository (NITRC-IR) [9], Federal Interagency Traumatic Brain Injury Research (FITBIR) [10], OpenNeuro [11], and National Institute of Mental Health Data Archive (NDA) [12] are dedicated to collecting neuro and brain imaging. The Cancer Imaging Archive (TCIA) [13] and Imaging Data Commons (IDC) [14] are cancer imaging repertories with TCIA provides images locally and IDC affords collections in the Cancer Research Data Commons (CRDC) cloud environment. Regarding breast cancer, the Cancer Research United Kingdom (UK) funds the OPTIMAM Mammography Image Database (OMI-DB) [15], and the University of Porto in Portugal funds the Breast Cancer Digital Repository (BCDR) [16], providing annotated breast cancer images and clinical details. Most of these repositories support data de-identification and quality control, except NITRC-IR and IDC. Additionally, some universities or institutions provide open-source datasets, such as Open Access Series of Imaging Studies (OASIS) [17], EchoNet-Dynamic [18], Cardiac Acquisitions for Multi-structure Ultrasound Segmentation (CAMUS) project [19], Chest X-ray [20] and Structured Analysis of the Retina (STARE) [21]. In China, the Huazhong University of Science and Technology provides an open resource named integrative CT images and CFs for COVID-19 (iCTCF) [22], which includes computed tomography (CT) images and clinical features of patients with pneumonia (including COVID-19 pneumonia). There is still a lack of databases that are dedicated to storing and accepting submissions for various diseases and modalities.

To address this issue, we established the Open Biomedical Imaging Archive (OBIA; https://ngdc.cncb.ac.cn/obia), a repository for archiving biomedical imaging data and related clinical data. As a core database resource in the National Genomics Data Center (NGDC) [23], part of the China National Center for Bioinformation (CNCB; https://ngdc.cncb.ac.cn/), OBIA accepts image submissions from all over the world and provides free open access to all publicly available data to support global research activities. OBIA supports de-identification, management, and quality control of imaging data, providing data services such as browsing, retrieval, and downloading, thus promoting the reuse of existing imaging data and clinical data.

## Implementation

### Database construction

OBIA is implemented using Spring Boot (a framework easy to create standalone Java applications; https://spring.io/projects/spring-boot) as the backend framework. The frontend user interface is developed using Vue.js (an approachable, performant and versatile framework for building web user interfaces; https://vuejs.org/) and Element UI (a Vue 2.0 based component library for developers, designers and product managers; https://element.eleme.cn/). The charts on the web page are constructed using ECharts (an open source JavaScript visualization library; https://echarts.apache.org). All metadata information is stored in MySQL (a free and popular relational database management system; https://www.mysql.com/).

### Image retrieval

Deep learning-based methods, such as scale-invariant feature transform (SIFT) [24], local binary patterns (LBP) [25], and histogram of oriented gradient (HOG) [26], demonstrate better performance compared to traditional methods. Deep neural networks excel at extracting superior features for retrieving multimodal medical images of various body organs [27] and enhancing ranking performance in the case of small sample sizes [28].

In OBIA, we leveraged TCIA multi-modal cancer data and utilized EfficientNet [29] as the feature extractor. We trained the model using a triplet network and an attention module to compress the image into a discrete hash value (**Figure 1**). We subsequently convert the trained model into TensorRT format to accelerate inference performance and reduce inference latency. We converted the trained model into TensorRT format to enhance inference performance and reduce latency. To store the hash codes, we employed Faiss, a high-performance similarity search library developed by Facebook AI Research, commonly used in deep learning. We computed image similarity using the hamming distance and returned the most similar images to the query. Our model achieved a mean average precision (MAP) value that surpassed the performance of existing advanced image retrieval models on the TCIA dataset.

**Figure 1.**
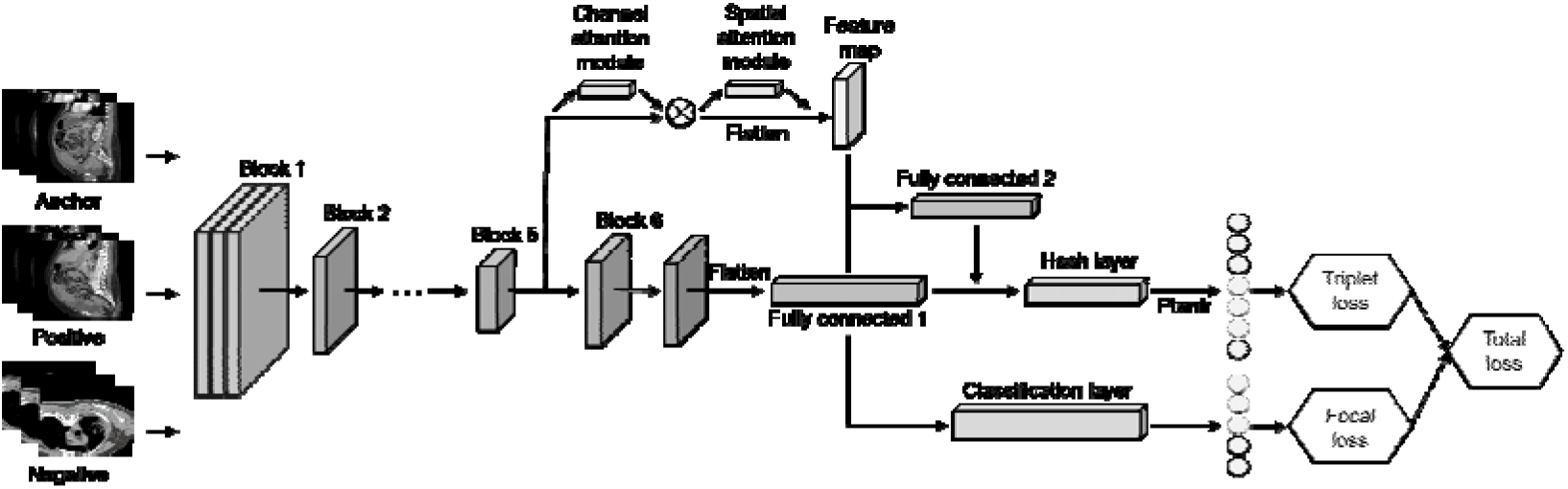
Deep triplet hashing based on attention and layer fusion module. The model uses EfficientNet-B6 as the backbone network and utilizes the CBAM attention module in Block5 to obtain feature maps. Layer fusion is employed in the fully connected layers, and focal loss and triplet loss are used to generate hash code and class embedding. CBAM, convolutional block attention module.

## Database content and usage

### Data model

Imaging data in OBIA are organized into five objects: Collection, Individual, Study, Series, and Image (**Figure 2**). “Collections”, bearing an accession number prefixed with “OBIA”, provides an overall description for a complete submission. “Individual”, possessing an accession number prefixed with “I”, defines the characteristics of a human or non-human organism receiving, or registered to receive, healthcare services. “Study”, adopting an accession number prefixed with “S”, contains descriptive information about radiological examinations performed on an individual. A study may be divided into one or more “Series” according to different logics, such as body part or orientation. “Image” describes the pixel data of a single Digital Imaging and Communications in Medicine (DICOM) file, and an image is related to a single series within a single study. Based on these standardized data objects, OBIA connects the image structure defined by the DICOM standard with actual research projects, realizing data sharing and exchange. Besides, each collection in OBIA is linked to BioProject [30] (https://ngdc.cncb.ac.cn/bioproject/) to provide descriptive metadata about the research project. And the individual in OBIA can be associated with GSA-Human [31] (https://ngdc.cncb.ac.cn/gsa-human/) by individual accession number, if available, which links imaging data with genomic data for researchers to perform multi-omics analysis.

**Figure 2.**
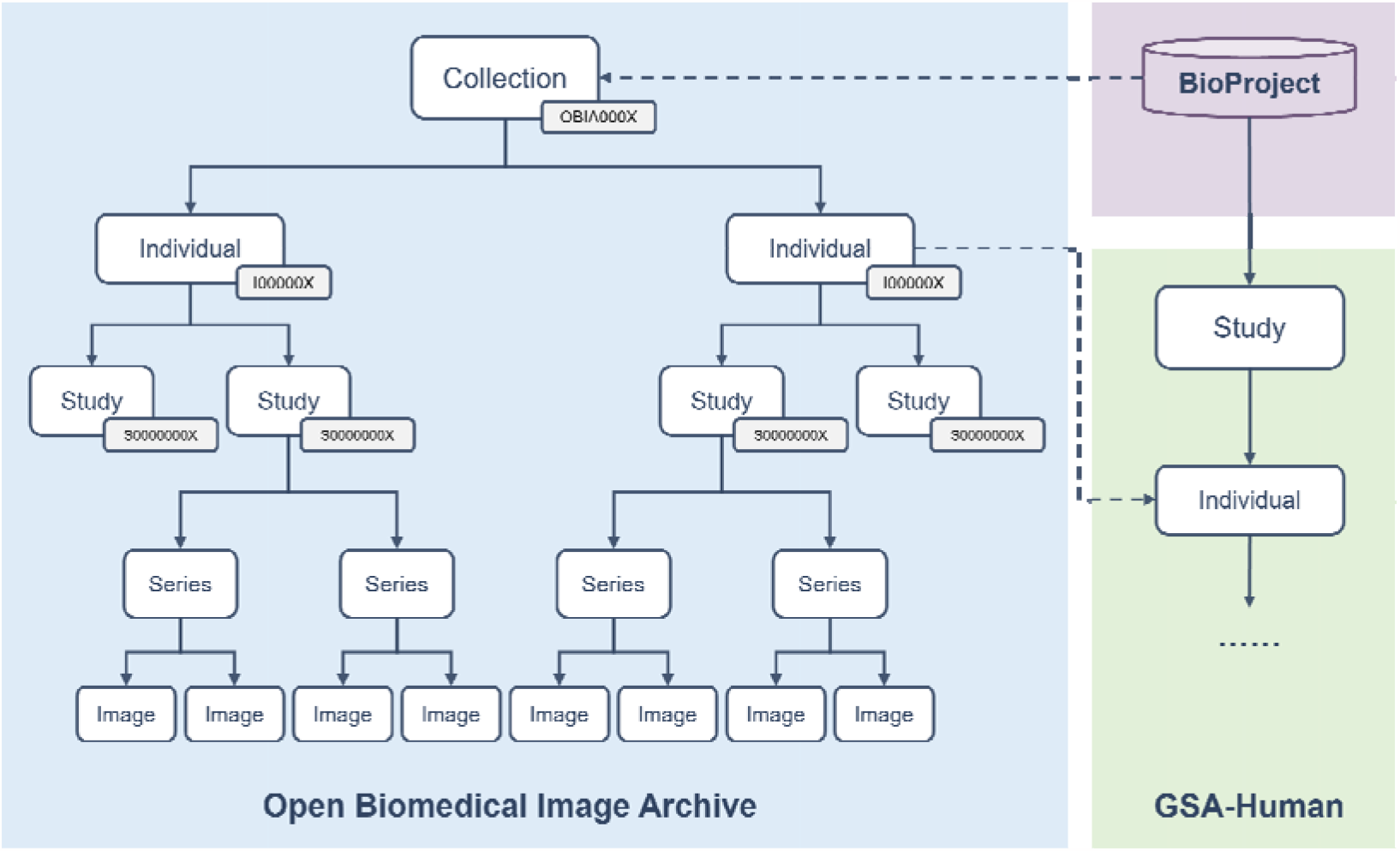
OBIA data model. The collection and individual in OBIA can be linked to BioProject and GSA-Human respectively. The accession numbers for data objects, including Collection, Individual, and Study, are indicated in the grey box. Collection accession numbers have a prefix of “OBIA” followed by four consecutive digits, Individual accession numbers have a prefix of “I” followed by six consecutive digits, and Study accession numbers have a prefix of “S” followed by eight consecutive digits.

### De-identification and quality control

Images may contain protected health information (PHI), and require appropriate handling to minimize the risk of patient privacy breaches. In order to retain as much valuable scientific information as possible while removing PHI, OBIA provides a unified de-identification and quality control mechanism (**Figure 3**) based on the DICOM PS 3.15 Appendix E: Attribute Confidentiality Profile (https://www.dicomstandard.org/). The key elements and rules we adopted include: 1) clean pixel data, 2) clean descriptors, 3) retain longitudinal temporal information modified dates, 4) retain patient characteristics, 5) retain device identity, 6) retain safe private tags.

**Figure 3.**
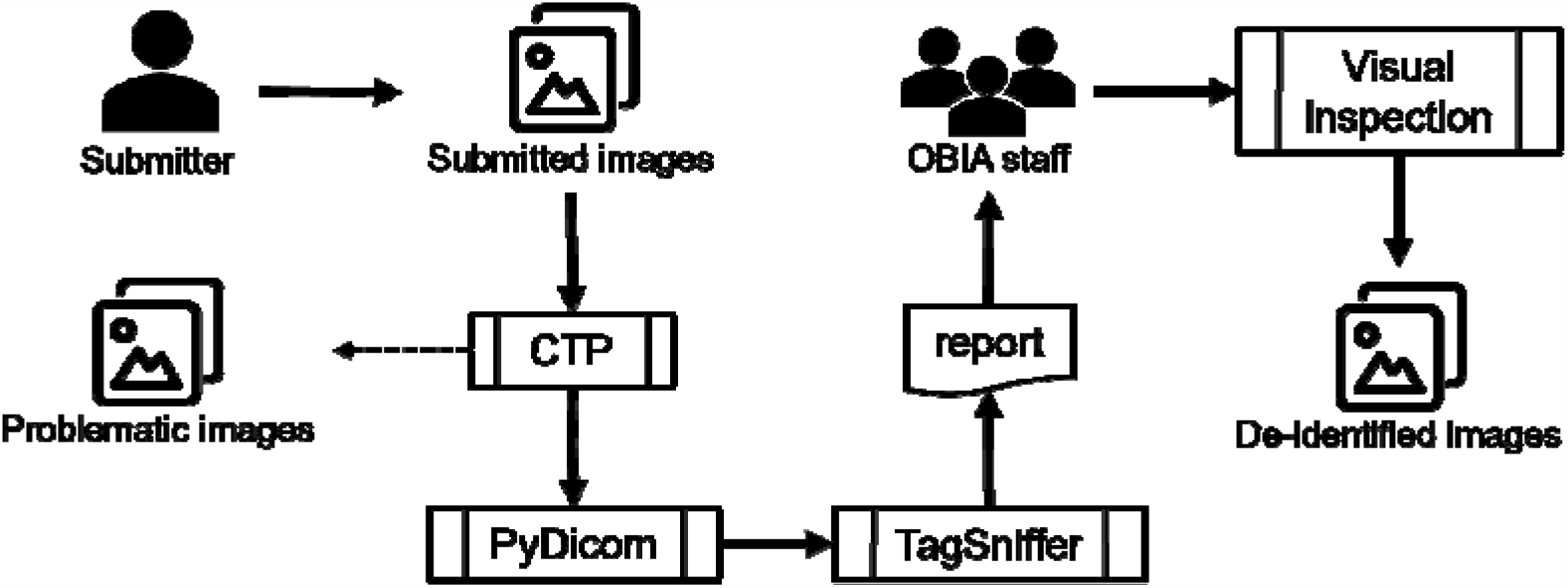
OBIA de-identification and quality control mechanism. Flowchart shows image submission, de-identification, and quality control steps. The de-identification steps include using CTP to process standard tags and PyDicom to handle private tags, and the problematic images will be isolated. The quality control steps include review of reports generated by TagSniffer and visual inspection of image pixels by OBIA staff. CTP, clinical trial processor.

OBIA utilizes the Radiological Society of North America (RSNA) MIRC clinical trial processor (CTP) (https://mircwiki.rsna.org/index.php?title=MIRC_CTP) for most of the de-identification work. We constructed a CTP pipeline and developed a common base de-identification script to remove or blank certain standard tags containing or potentially containing PHI. This script also maps local patient IDs to OBIA individual accessions. As for private tags, since they are vendor-specific and the scanner-specific and can contain almost anything, we use PyDicom (https://pypi.org/project/pydicom/) to retain attributes that are purely numeric. Some studies determine private element definitions by reading manufacturers’ DICOM conformance statements [32]. However, in our practice, the wide variety of image sources made this work too time-consuming, and some conformance statements were not available. In addition to metadata elements, ultrasounds or screen captures, usually add some “burned in” annotations in the pixels data to interpret the images, which may also contain PHI. We provide a filter stage to identify these images.

After the de-identification process is completed, OBIA runs a quality control procedure. Some problematic images are isolated, such as images with a blank header or missing patient ID, corrupted images, other patient images mixed in, etc. Submitters can provide relevant information to repair the images or discard them entirely. Duplicate images are removed, leaving only one. Then we use TagSniffer (https://github.com/stlstevemoore/dicom-tag-sniffer) to generate a report for all images. All DICOM elements in the report are carefully reviewed to ensure that they are free of PHI, and certain values (e.g., patient ID, study date) are modified as expected. In addition, we perform a visual inspection of each image series to ensure that no PHI is contained in the pixel values and that the images are visible and uncorrupted.

### Data browse and retrieval

OBIA provides user-friendly web interfaces for data query and browsing. Users can browse data of interest by specifying non-image information (e.g., age, gender, disease) and/or the imaging data extracted from the DICOM header (e.g., modality, anatomical site). Users can also search for data by entering the accession number. OBIA allows users to browse the basic information of collection (title, description, keywords, submitter, data accessibility, etc.), individual, study (study description, study age, study date, etc.), series (modality, anatomical site, series description, etc.) and view thumbnails of images.

OBIA also provides image retrieval functionality designed to find images similar to a query. Users can use this function by clicking the “Image Retrieval” button on the homepage. After uploading the image, the image retrieval model employs the hamming distance to return the top 30 images that are closest to the queried image, serving as the nearest neighbor images. Users have the option to click on these returned images and review their respective image metadata.

### Data access and download

OBIA states that the data access policies are set by data submitters. There are two different types of data accessibility: open access and controlled access. Open access means that all data is public for global researchers if released, whereas controlled access means that data can be downloadable only after being authorized by the submitter. OBIA supports online data requests and reviews. Before applying, users need to register and log into OBIA via the Beijing Institute of Genomics (BIG) single sign-on (SSO; https://ngdc.cncb.ac.cn/sso/) system. Applicants shall provide their basic information, specify the scope of data usage, and promise not to attempt to restore the privacy information in the data. The download link will only be provided to the applicant if the data owner approves the request.

### Data submission

OBIA accepts submissions of biomedical imaging data in DICOM format from clinical or specific research projects. To create a submission, users need to log in, fill in the basic information of the collection, and email the necessary clinical information. Image data will be transferred to OBIA offline after de-identification. Users can either use the de-identification process recommended by OBIA or apply their own methods to de-identify the image. Quality control will be conducted for all the data. OBIA will assign unique accession number to each collection, individual, and study and arrange images into a standard organization. In order to ensure the security of submitted data, backup copies will be stored on physically separate disks. Finally, metadata will be extracted from the image file headers and stored in the database to support data queries.

### Data statistics

As of September 2023, OBIA has housed a total of 937 individuals, 4,136 studies, 24,701 series, and 1,938,309 images, covering 9 modalities and 30 anatomical sites. Representative imaging modalities are computed tomography (CT), magnetic resonance (MR), and digital radiography (DX) (Table S1). Anatomical sites include abdomen, breast, chest, head, liver, pelvis, and so on. (Table S2). The first batch of collections submitted to OBIA came from the Chinese People’s Liberation Army (PLA) General Hospital, including imaging data of three major gynecological tumors: endometrial cancer, ovarian cancer, and cervical cancer. The data were divided into four collections, and **Table 1** shows the number of individual, study, series, and image of each collection. In addition, we collected associated clinical metadata, such as demographic data, medical history, family history, diagnosis, pathological types, and treatment methods.

**Table 1.** Number of Individual, Study, Series, and Image of each Collection.

| Accession | Disease | Individual | Study | Series | Image |
| --- | --- | --- | --- | --- | --- |
| OBIA0001 | Endometrial Cancer | 316 | 1587 | 8941 | 779,171 |
| OBIA0002 | Ovarian Cancer | 202 | 1548 | 7151 | 548,174 |
| OBIA0003 | Cervical Cancer | 117 | 675 | 3943 | 307,101 |
| OBIA0004 | Endometrial Cancer | 302 | 326 | 4666 | 303,863 |

*Note:* The data statistics are up to September 2023.

### Perspectives and concluding remarks

OBIA is a centralized repository of de-identified biomedical imaging data. Different from existing related databases, OBIA is characterized by publishing imaging data and clinical information from a wide range of imaging modalities in a common DICOM format. Algorithm developers can convert and format as needed, and clinicians and researchers can combine clinical data with images for further analysis. Unlike the Health Insurance Portability and Accountability Act (HIPAA) in the United States and General Data Protection Regulation (GDPR) in Europe, China has Personal Information Protection Law (PIPL), Data Security Law, and other regulations for medical records and health information, which include elements similar to HIPAA and GDPR. OBIA strictly follows Chinese legal regulations for data processing, quality control and data sharing, especially for removing PHI. It also promotes data sharing among data submitters and users while ensuring compliance with these laws. As a core database resource within NGDC, OBIA is seamlessly integrated with BioProject and GSA-Human, facilitating the harmonious integration of imaging and genomic data to support multi-omics analysis. In essence, OBIA serves as a dependable platform for sharing clinically relevant imaging data among researchers from diverse institutions, effectively bridging the gap within China’s biomedical imaging database landscape.

In the future, we will continue to upgrade the infrastructure of OBIA and increase security protection measures to realize long-term secure storage, management and access to a large volume of images. At the same time, we will collect more types of biomedical imaging data and gradually increase the corresponding genomic data to expand our data resources. To facilitate data submission and ensure privacy security, we will further optimize the image de-identification process and explore the use of machine learning-based optical character recognition (OCR) [33] method to remove PHI from image pixels. We will also improve quality control and background automatic review processes to speed up data submission. In compliance with applicable regulations and ethical norms, OBIA’s goal is to preserve as much effective image metadata as possible to provide researchers with high-quality imaging data. Furthermore, we plan to develop more intuitive and interactive web interfaces according to users’ needs, increase database functions, and integrate related online tools to help analyze biomedical images. In addition, we intend to optimize the image retrieval model to offer users more convenient and precise image retrieval services. Finally, we call for collaborators to collectively build OBIA, submit image data, break down data silos, catalyze new biomedical discoveries, and provide the possibility to create personalized treatments.

## Ethical statement

The collection of human imaging data was approved by the Local Ethical Committees in the First Medical Center of the Chinese PLA General Hospital (Approval No. S2022-403). The written informed consent was obtained from the participating subjects.

## Data availability

OBIA is publicly available at https://ngdc.cncb.ac.cn/obia.

## CRediT author statement

**Enhui Jin:** Investigation, Methodology, Software, Writing – original draft. **Dongli Zhao:** Resources. **Gangao Wu:** Investigation, Methodology, Software, Writing – original draft. **Junwei Zhu:** Software. **Zhonghuang Wang:** Methodology. **Zhiyao Wei:** Resources. **Sisi Zhang:** Methodology. **Anke Wang:** Software, Writing – review & editing. **Bixia Tang:** Resources. **Xu Chen:** Resources. **Yanling Sun:** Investigation, Methodology, Writing – review & editing, Project administration. **Zhe Zhang:** Investigation, Methodology, Writing – review & editing. **Wenming Zhao:** Conceptualization, Methodology, Writing – review & editing, Supervision, Funding acquisition. **Yuanguang Meng:** Conceptualization, Methodology, Writing – review & editing. All authors read and approved the final manuscript.

## Acknowledgments

This work was supported by grants from the Strategic Priority Research Program of the Chinese Academy of Sciences (Grant No. XDB38050300), Genomics Data Center Operation and Maintenance of Chinese Academy of Sciences (Grant No. CAS-WX2022SDC-XK05), and the Key Technology Talent Program of the Chinese Academy of Sciences.

## Supplementary material

**Table S1 Number of Individual, Study, Series, and Image of each imaging modality**

**Table S2 Number of Individual, Study, Series, and Image of each anatomical site**

