## Supplementary material for "OBIA: An Open Biomedical Imaging Archive": Table S1

**Table S1** **Number of Individual, Study, Series, and Image of each imaging modality**

| **Modality** | **Individual** | **Study** | **Series** | **Image** |
| --- | --- | --- | --- | --- |
| CR | 327 | 603 | 936 | 936 |
| CT | 453 | 1608 | 8739 | 1,200,188 |
| DX | 433 | 1059 | 2913 | 3072 |
| IO | 14 | 23 | 28 | 28 |
| MG | 4 | 4 | 4 | 16 |
| MR | 597 | 831 | 12,036 | 734,024 |
| PX | 2 | 2 | 2 | 2 |
| RF | 4 | 4 | 35 | 35 |
| XA | 1 | 8 | 8 | 8 |

*Note:* The data statistics are up to September 2023. CR, computed radiography; CT, computed tomography; DX, digital radiography; IO, intra-oral radiography; MG, mammography; MR, magnetic resonance; PX, panoramic x-ray; RF, radio fluoroscopy; XA, x-ray angiography.
