## Supplementary material for "OBIA: An Open Biomedical Imaging Archive": Table S2

**Table S2 Number of Individual, Study, Series, and Image of each anatomical site**

| **Anatomical site** | **Individual** | **Study** | **Series** | **Image** |
| --- | --- | --- | --- | --- |
| Abdomen | 229 | 540 | 2848 | 336,538 |
| Ankle | 2 | 2 | 4 | 4 |
| Aorta | 1 | 1 | 2 | 2354 |
| Brain | 1 | 1 | 11 | 245 |
| Breast | 4 | 5 | 10 | 2364 |
| Chest | 603 | 1970 | 5827 | 345,434 |
| C-spine | 13 | 13 | 19 | 124 |
| Extremity | 4 | 4 | 15 | 5804 |
| Foot | 2 | 2 | 4 | 4 |
| Hand | 2 | 2 | 3 | 3 |
| Head | 33 | 42 | 176 | 6532 |
| Heart | 6 | 6 | 21 | 1099 |
| Hip | 2 | 3 | 24 | 1812 |
| Jaw | 9 | 15 | 20 | 20 |
| Knee | 9 | 13 | 44 | 1465 |
| Leg | 1 | 2 | 3 | 3 |
| Liver | 16 | 26 | 261 | 21,509 |
| L-spine | 12 | 12 | 28 | 92 |
| Lung | 1 | 1 | 5 | 761 |
| Neck | 4 | 9 | 44 | 5583 |
| Pelvis | 416 | 615 | 8501 | 517,962 |
| Prostate | 3 | 3 | 36 | 3480 |
| Shoulder | 4 | 7 | 31 | 488 |
| Skull | 1 | 1 | 1 | 1 |
| Spine | 12 | 16 | 74 | 2339 |
| Thorax | 2 | 2 | 3 | 3 |
| T-spine | 1 | 1 | 9 | 144 |
| Unknown | 362 | 896 | 4002 | 483,895 |
| Upper limb | 1 | 1 | 2 | 2 |
| Wrist | 1 | 1 | 1 | 2 |

*Note:* “Unknown” means that there is no information available in the DICOM tag “body part examined”. The data statistics are up to September 2023. DICOM, digital imaging and communications in medicine.
